# Velcro-like mannose and slime-like sialic acid interactions guide self-adhesion and aggregation of virus N-glycan shields

**DOI:** 10.1101/2020.05.01.072769

**Authors:** Eric Ogharandukun, Wintana Tewolde, Elbethel Damtae, Songping Wang, Andrey Ivanov, Namita Kumari, Sergei Nekhai, Preethi L. Chandran

**Author notes:** Corresponding author, Preethi L. Chandran, PhD, Associate Professor, Department of Chemical Engineering, College of Engineering and Architecture, Howard University, Department of Biochemistry and Molecular Biology, College of Medicine, Howard University, Address: 1011 LK Downing Hall, 2300 6th Street, NW, Howard University, Washington, DC 20059. **Email:**.

## Abstract

The surfaces of cells and pathogens are covered with short polymers of sugars known as glycans. Complex N-glycans have a core of three mannose sugars, with distal repeats of N-acetylglucosamine and galactose sugars terminating with sialic acid (SA). Long-range slime-like and short-range Velcro-like self-adhesions were observed between SA and mannose residues, respectively, in ill-defined monolayers. We investigated if and how these adhesions translate when SA and mannose residues are presented in complex N-glycan shields on two pseudo-typed viruses brought together in force spectroscopy (FS). Slime-like adhesions were observed between the shields at higher ramp rates, whereas Velcro-like adhesions were observed at lower rates. The complex glycan shield appears penetrable at the lower ramp rates allowing the adhesion from the mannose core to be accessed; whereas the whole virus appears compressed at higher rates permitting only surface SA adhesions to be sampled. The slime-like and velcro-like adhesions were lost when SA and mannose, respectively, were cleaved with glycosidases. While virus self-adhesion in FS was modulated by glycan penetrability, virus self-aggregation in solution was only determined by the surface sugar. Mannose-terminal viruses self-aggregated in solution, while SA-terminal ones required Ca^2+^ ions to self-aggregate. Viruses with galactose or N-acetylglucosamine surfaces did not self-aggregate, irrespective of whether or not a mannose core was present below the N-acetylglucosamine surface. Well-defined rules appear to govern the self-adhesion and -aggregation of N-glycosylated surfaces, regardless of whether the sugars are presented in ill-defined monolayer, or N-glycan, or even polymer architecture.

## 1. Introduction

The surface of eukaryotic cells and pathogens is covered with short, branched sequences of sugars known as glycans protruding from the proteins and lipids within the membrane.(1–5) More than 70% of these are N-glycans characteristically attached via the amine groups on proteins and having a core of two N-acetyl glucosamine (GlcNAc) followed by three mannose (man) sugars (Fig. 1). There are three types of N-glycans depending on the sequence of the sugar branches distal to the core: (i) high mannose type where all branches have mannose sugars, (ii) complex type where branches have repeat sequences of GlcNAc and galactose residues capped by sialic acid (SA), and (iii) hybrid type where one branch is high-mannose type and the other branch is complex type.(6–8) Fucose sugars may be attached to GlcNAc in the glycan core.(9)

**Fig.1:**
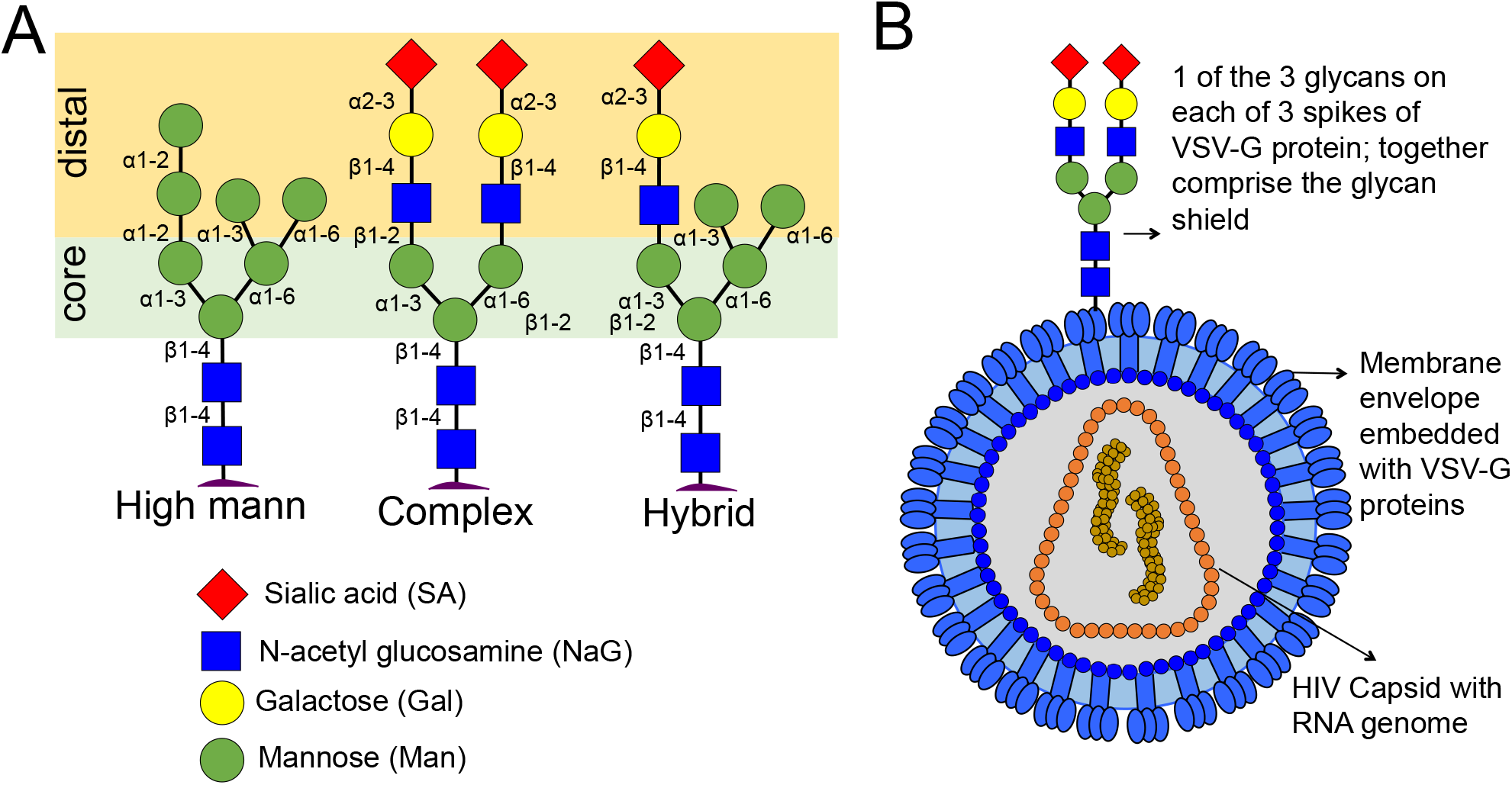
[A] Types of N-glycans protruding from proteins in cell membranes. [B] Presentation of complex N-glycans within the glycan shields of HIV-1 virus pseudotyped with VSV-G proteins.

The protein binding of glycan sugars is a key event in the communication cascade between cells, cells and environment, and host and pathogens.(10–12) Debilitating malfunction of organ systems occurs in congenital disorders of glycosylation, a condition where glycan processing proteins are defective, indicating the criticality of glycans for biological function.(13–15) In addition to presenting sugars in conformations recognized by proteins, the branched architecture of N-glycans was also found critical for post-natal survival, protection against autoimmunity, and development of the nervous system.(16–18) The branching of N-glycans enhances biochemical signals and increases during tumor metastasis and apoptosis.(19) The multiplicity of galactose presentation, for instance, recruits clusters of galactose-binding lectins (galectins) close to the cell membrane, which crosslink neighboring membrane domains.(20–22) Reduced cell migration, growth arrest, and enhanced auto immunity are observed as a result. In sequence of sugars also plays a role; each sugar in N-glycan sequence can participate in mutually-exclusive biochemical pathways.(23) While the role of N-glycan presentation in protein-glycan binding and signalling is increasingly appreciated; the biophysical interactions resulting from the N-glycan presentation, with its sequence and architectural placement of sugars, remains to be investigated.

In single-sugar monolayer studies, two types of adhesions were observed between the sugars present in N-glycans. Perera et al.(24) reported self-latching adhesions between monolayers of mannose (man). These man-man adhesions can be described as Velcro-like; they released short-range with a major peak at the separation interface between the layers. The Velcro-like releases also occurred when one of the mannose layers was present under tiers of non-mannose sugars in complex N-glycan shields, suggesting that the latter did not impede access to the mannose core or cross-adhere to mannose.(25) SA was the only other N-glycan sugar to exhibit self-adhesion in monolayer studies, however its adhesion release was more slime-like, occurring over long distances with multiple unabating peaks (saw-tooth pattern) and mediated by salt. In solution, mannose-nanoparticles coated, whereas SA-nanoparticles aggregated virus with complex N glycan shields.

Due to the their conserved composition and preponderance on biological surfaces, any rules relating the biophysics and composition of N-glycans will have wide implications for predicting adhesion and aggregation patterns in cell-cell, cell-pathogen, and pathogen-pathogen contacts. However, above studies were performed with artificial monolayers of sugars generated by inserting linker molecules into the sugars. The sugars in these monolayers had random linkages and orientations, unlike in N-glycans where there is a defined branched architecture with accompanying sugars. It is not clear if the above observations with monolayers would apply when the same sugars are present in N-glycan context, and how adhesions between single sugars translate for the multi-sugar tiered architecture of N-glycans.

The goal of the study is to determine how glycan sugar-sugar adhesions, observed previously with disordered monolayers, manifest in the ordered and biologically relevant presentation of N glycans. Human Immunodeficiency Virus 1 (HIV-1) exhibits dense glycan shields contributed predominantly by one type of membrane protein in the envelope membrane. VSV envelope protein (VSV-G) is a trimer with multiple complex N-glycans attached.(26) Shields of complex N-glycan shields can be generated by pseudo-typing (i.e., engineering to display a desired envelope protein) HIV-1 virus with VSV-G envelope proteins (Fig. 1B). Our study design involves bringing two complex N-glycan shields of virus into forced contact with force spectroscopy (FS) or into free contact by diffusion in solution. The changes in adhesion forces and aggregation pattern are tracked as layers of sugars are shaved off the glycan shield with site-specific glycosidases in order to determine the self-adhesion rules between N-glycan sugars, particularly mannose and SA, when in N-glycan architectural context.

## 2. Materials and Methods

### 2.1 Materials

293T cells and 293 cells were purchased from ATCC (Manassas, VA). The pHEF-VSVG expression vector (courtesy of Dr. Lung-Ji Chang) and pNL4-3.Luc.R^−^E^−^ (Courtesy of Dr. Nathaniel Landau) were obtained from the NIH AIDS Research and Reference Reagent Program. Glycosidases and lectins were purchased from Millipore Sigma Inc., St. Louis, MO. The glycosidase β-N-Acetylglucosaminidase (NAG) from *Canavalia* ensiformis was used to expose the mannose core. The enzyme β-mannosidase from *Helix pomatia* was used to remove the mannose core. Neuraminidase from *Clostridium perfringens* was used to expose galactose residues, and α-mannosidase from *Canavalia ensiformis* was used to cleave between the two tiers of mannose in the core. β-galactosidase (β-gal) from *Aspergilus* oryzae was used to remove galactose residues.

### 2.2 Production and characterization of virus

Vesicular stomatitis virus G protein (VSV-G)**-** pseudotyped pNL4-3.Luc.R-E-virus was prepared as previously described. (27) In brief, HEK293T cells were co-transfected with pNL4-3.Luc.R-E-^−^ HIV-1 genomic vector and pHEF VSV-G expression vector. The former does not express the HIV envelope protein Gp120, whereas the latter encodes the VSV-G envelope protein. Media with virus was collected at 72 hrs post transfection, concentrated, and stored at −70°C. In a previous study, the pseudotyped HIV-1 R^−^E^−^/VSV-G virus was found morphologically similar to wild type HIV-1 virus in that it contains a cone-shaped capsid within a membrane envelope, but the interactions of the glycan shield was consistent with the presence of complex N-glycans from VSV-G envelope proteins. (25)

### 2.3 Virus attachment to AFM probe

Virus was attached to a gold-coated probe using dithiobis succinimidyl propionate (DSP) linkers (Thermo Scientific). Dithiobis succinimidyl propionate (DSP) (4 mg) was dissolved in 1 ml dimethylsulfoxide (DMSO) and applied on gold-coated AFM probes for 30 minutes at room temperature. DMSO reduces the disulfide bond in the middle of the bi-functional linker. The released molecules assemble to the probe via thiol groups at one end and to display NHS ester groups on the free end. The probes were rinsed with water to activate the NHS groups. The HIV-1 R^−^E^−^/VSV-G virus in phosphate-buffered saline (PBS) was immediately added to the probe and incubated for 1 hr at room temperature to allow the activated NHS groups to react with amines on the virus surface. The probe was rinsed with PBS to remove all cross-linker products including NHS leaving groups and unconjugated virus, and used immediately for experiments. The grafting schematic is presented in Fig. 2. By assembling the linkers on the surface and then overlaying the virus, the latter’s surface probed in FS was not exposed to crosslinking reactants and organic solvents.

**Figure 2:**
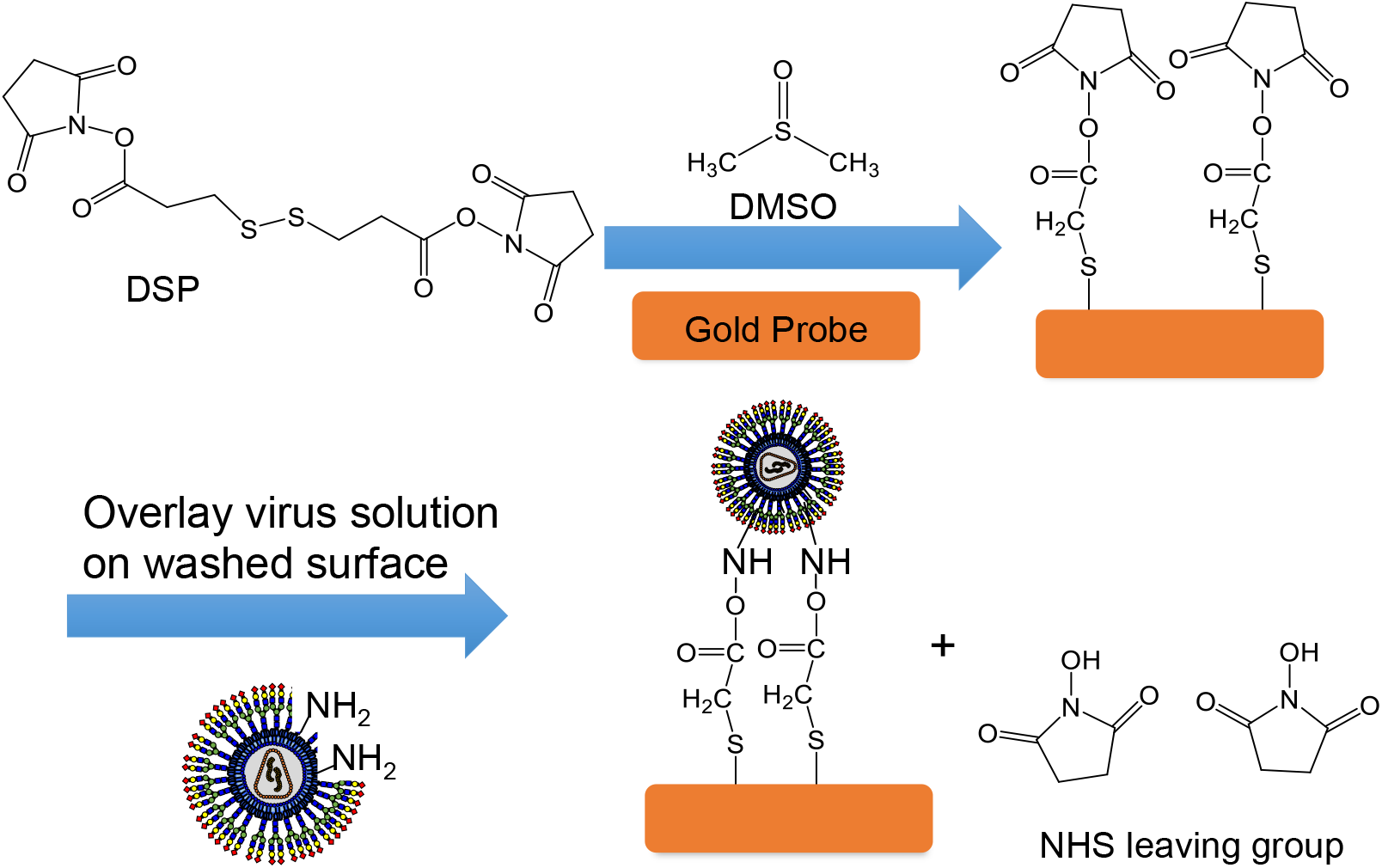
The reaction scheme for attaching virus to the gold surface.

### 2.4 Nanoindentation of HIV-1 pseudotyped virus attached to virus probe

The HIV-1 R-E-/VSV-G virus surface has negatively charged SA sugars which adsorbs on positive-charged APTES-coating on mica surface. The APTES coating was generated by placing 100μl of 5mM (3-aminopropyl) triethoxysilane (APTES) on freshly cleaved mica surface and incubating for 30 min, followed by washing with water and then PBS. HIV-1 R^−^E^−^/VSV-G (6μl in PBS) was applied to the APTES surface for about 8 minutes. Then the surface was washed, and covered with about 100 μl PBS before mounting onto the sample stage of a Multimode/PicoForce system (NS-V controller, Bruker Nano-surfaces Inc., Santa Barbara, CA). A fluid cell retained the buffer during indentation experiments. NPG-10 cantilevers (Bruker Inc., Billerica, MA) were functionalized with HIV-1 R^−^E^−^/VSV-G as described in Fig. 2 and mounted on the AFM cantilever holder. The spring constant (k) of the cantilever was determined by standard thermal tuning methods. The contact force for initiating cantilever retraction (i.e., maximum indentation force) was set at ~1nN and the ramp size at ~500nm. Force spectroscopy data was analyzed using in-house MATLAB code to quantify descriptive features of each curve. Indentation thickness is calculated as the difference between the separations where indentation force becomes higher than zero and where the maximum indentation force is reached (see Fig. 3B). Adhesion length is calculated as the separation span over which the adhesion releases. Adhesion energy is the work done to release an adhering AFM probe, and is calculated as the area under the negative adhesion force during retraction.

**Figure 3:**
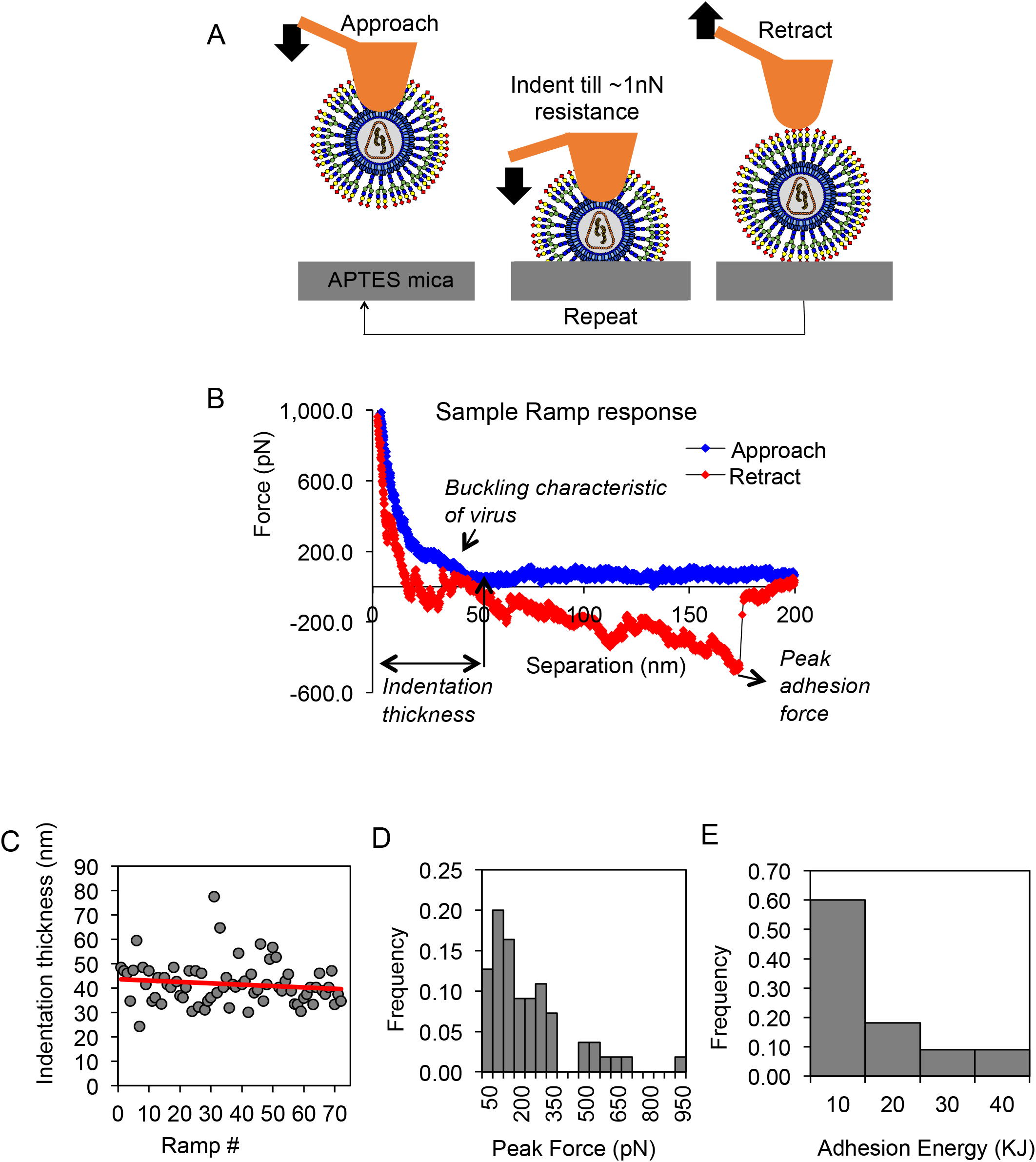
[A] Experimental scheme for testing resilience of virus attachment to AFM probe. The probe with attached virus is repeatedly pushed into opposite-charged APTES surface till 1nN is reached and retracted. [B] A typical force curve observed when probe-virus is compressed against APTES surface has buckling features in approach and adhesion to surface in retraction. [C] Evolution of indentation thickness, a measure of virus compliance, as the probe-virus is repeatedly ramped into the APTES surface. [D] Range of peak adhesion forces with opposite-charged APTES surface endured by the virus during repeated ramping and retraction. [E] Range of adhesion energies resisting probe-virus pull-out from the APTES during course of repeated ramping.

### 2.5 Cleaving sugars on bound and free virus

For force spectroscopy, cleavage was performed on HIV-1 R^-^E^-^/VSV-G attached to AFM probes or those adsorbed on APTES mica surfaces. The virus surface was rinsed with 100 μL PBS. 60μL of 1U/ml glycosidase and 2.5 mg/mL BSA in PBS was overlaid on the virus surface. 45 min to 1-hr incubation was found sufficient to cleave sugar tiers without leaving significant interaction traces from uncleaved sugars. The enzyme mixture and cleaved sugars were washed from the virus surface using PBS. Since sugar cleavage was performed on surface-bound virus, cleaved products and enzymes could be removed by simple washing of the surface. Moreover the structural stability of the virus would also be retained to some extent. For cleaving virus in solution, stock enzyme solutions were prepared as per manufacturer recommendations and mixed with virus to obtain final concentrations of ~0.15 U/ml. The completion of cleavage was monitored from virus aggregation with DLS and by increasing the enzyme concentration. Virus aggregate sizes observed in DLS stabilized by 1 hour for the concentrations used in this study.

### 2.6 Dynamic Light Scattering (DLS)

DLS measurements were performed in 40 μL disposable cuvettes with a Malvern Zetasizer ZS (Malvern Instruments, Inc., Westborough, MA) as described previously.(25) Hydrodynamic diameters D_H_ were extracted from the correlation curves using an inhouse Matlab software based on the Sequential Extraction of Late Exponentials (SELE) algorithm. (28)

## 3. Results

### 3.1 Confirming virus attachment to AFM probe

NHS-terminated linkers were assembled on gold-coated AFM probes via chemisorption of thiol group at the other end. The NHS group traps overlaid virus by reacting with amines on membrane proteins. Since the virus is ~ 80 nm in diameter, i.e., larger than the spacing between linkers in assembled monolayers, each particle is expected to bind to the gold surface of an AFM probe with several short linkers. To verify if the virus remains structurally intact through the attachment procedure and can survive the repeated push and pulls in force spectroscopy experiments, the following test was performed.

The probe-attached virus was repeatedly pushed into APTES-coated mica until 1nN of indentation force was reached and withdrawn (Fig. 3A). When the probe approaches the APTES surface (i.e., *separation* between the probe and surface decreases), the indentation *force* rises as the virus compresses against the surface (blue curve in Fig. 3B). The force rise is not steady, however, but flattens for several nms before rising again. A leveling mid-way in the indentation force typically signifies buckling-like deformations in shell and micelle materials, indicating that the membrane and capsid structure in the probe-bound virus are physically intact. The resistance profile of the probe-bound virus being pushed into a surface is similar to that observed when surface-adsorbed virus was indented by a bare probe (25), confirming that virus was attached to the AFM probe without significant alterations in structure.

Indentation thickness is the separation or depth the virus is compressed by before 1.2 nN of resistance is registered in the approach curve. The indentation thickness would drop to zero if virus gets dislodged from the probe or completely squished. Over the course of repeated pounding into the APTES surface (~90 ramps shown in Fig. 3C), the indentation thickness of the virus remained around 40 nm. The overall consistency in indentation thickness confirms that the probe-bound virus was able to sustain repeated ramping without detaching, rupturing, or collapse. The indentation thickness showed a modest downward trend with ramp number (Fig. 3C), suggesting that in this experiment the continuous pounding could be irreversibly compressing or stiffening the virus.

When the virus-bound probe was retracted after compressing into the APTES surface, a negative adhesion force was observed (Fig. 3B, red curve), indicating that the positively charged APTES surface was resisting the virus pull out. The distribution of adhesion forces and energies endured by the virus during the course of the repeated ramping are summarized in Fig. 3D. Since the indentation thickness did not shift to zero (Fig. 3C), the attachment between the probe and virus can survive the 0.9 nN of force and 40 KJ of adhesion energy encountered in this experiment without detaching. In cases where virus was pushed into the APTES surface with greater indentation force of 2nN, greater adhesion forces were registered during retraction, without dislodging or disrupting the virus on the probe (data not shown). From the range of adhesion forces survived in these experiments, the attachment between virus and probe can be considered stronger than the adhesion forces to be sampled between glycans in subsequent experiments.

### 3.2 Rate-dependent shift from slime- to velcro-like self-adhesion between complex N-glycan shields

We tested the adhesion between the complex N-glycan shields on two viruses, one bound to probe and the other adsorbed on APTES (Fig. 4A). The viruses were brought into contact until an indentation force of 1nN was registered, so that only the interactions between glycan shields were sampled (25). As the two virus glycan shields retracted after contact, the nature of adhesion release changed with the indentation rate. At the lower rates (toward 50 nm/sec), the release occurred in one predominant peak around the point of pullout between the two viruses (Fig. 4B). Such short-range ‘brittle’ or Velcro-like adhesion releases were observed earlier between two monolayers of mannose residues.(24) At the higher indentation rates (towards 500 nm/sec), however, the adhesion released in multiple peaks increasing in magnitude even far from the contact point between the two surfaces (Fig. 4D). Such long-range sawtooth-like releases were observed earlier between two SA monolayers, and can be characterized as ‘slime-like’ (releasing in spurts over long range). In between the higher and lower rates, the adhesion release was usually a mix of the two types (Fig. 4C).

**Figure 4:**
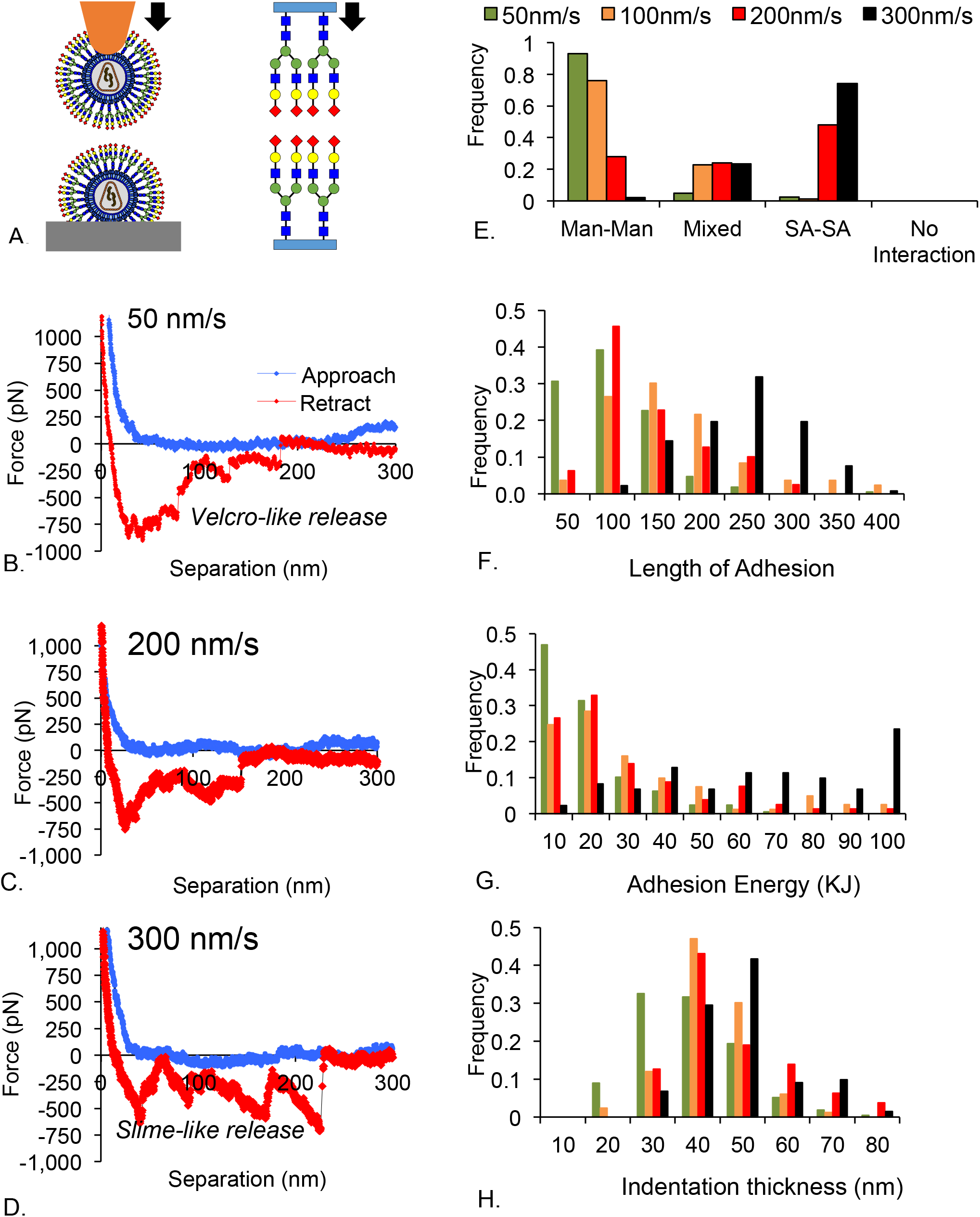
SA-SA and man-man adhesions appear at high and low rates, respectively, of virus-virus interaction. [A] Experimental setup where two viruses are brought into contact upto a force of 1.2nN and retracted. [B – D] Forces representative of velcro-like, mixed, and slime-like adhesions were predominantly observed at low (50nm/sec), intermediate, and high (>300nm/sec) rates. Velcro-like adhesions are characterized by a single dominant peak close to the contact point between the two surface, and were previously observed between man-man monolayers. Slime-like adhesions are distinguished by a saw-tooth like profile of release with several dominant peaks which continue undiminished away from the contact point between the two surfaces, and were previously observed in SA-SA monolayers. Mixed type has features of both. [E] The frequency of occurrence of the three adhesion types depends on the indentation rate, with slime-like adhesions occuring toward the higher rates and velcro-like adhesions occuring at lower rates. [F] The distribution of adhesion lengths measured during retraction is dependent on the indentation rate, with the longer lengths occuring towards the higher rates. [G] The distribution of adhesion energies measured during retraction is dependent on the indentation rate, with higher energies occuring towards the higher rates. [H] The distribution of indentation lengths was not dependent on indentation rate, indicating similar virus populations being sampled at all rates. (n > 65 for each ramp rate, 3 samples and 2 runs).

In >500 curves (multiple viruses, adsorption surfaces, and stocks) analyzed per indentation rate, the ‘slime-like’ adhesions occurred predominantly at higher ramp rates (above 300nm/sec) whereas ‘Velcro-like’ adhesions were frequent at the lower ramp rates (Fig. 4E). At intermediate rates, adhesion curves with both features (‘mixed’) were likely to occur. There were no instances where the two viruses retracted without adhesion.

Quantifiable features of the force curves like adhesion length and adhesion energy also exhibited rate-dependent trends (Fig. 4F,G). The adhesion lengths clustered around 100 nm at the higher indentation rates (consistent with long-range slime-like release) and around 20 nm at the lower rates (consistent with short-release Velcro-like release at these rates) (Fig. 4F). The adhesion energy distributed around larger values at the higher indentation rates (consistent with long-range release), and shifted to lower values at the lower rates (consistent with short-range Velcro-like adhesions) (Fig. 4G). The indentation thickness, however, did not change between the different rates (Fig. 4H), indicating that a similar population of viruses was sampled at all the rates.

### 3.3 Rate-dependent shift in adhesion pattern relates to penetrability of complex N-glycan shield

VSV-G glycoproteins are known to predominantly have complex N-glycans and therefore present a glycan shield with surface SA residues and core mannose residues. Accordingly, we expected to observe SA-SA slime-like adhesions occurring between two complex glycan shields at all rates. Instead man-man like Velcro adhesions occurred at low rates of indentation, and SA-like adhesions at only the higher rates. To determine the cause of the rate-dependent shift, we first needed to verify if there were surface mannose sugars on the virus in the regions being sampled at the lower indentation rates.The same spot on the glycan shield of two viruses was approached and retracted at multiple rates. The velcro-like release at lower rates shifted to slime-like sawtooth release at higher rates even when at the same location (Fig. 5), confirming that the rate-dependence shift was not due to spatial variations in glycan types. Lectin binding studies in solution (Sec. 3.6) also verified that mannose residues were not present terminally in the virus glycan shield.

**Fig.5.**
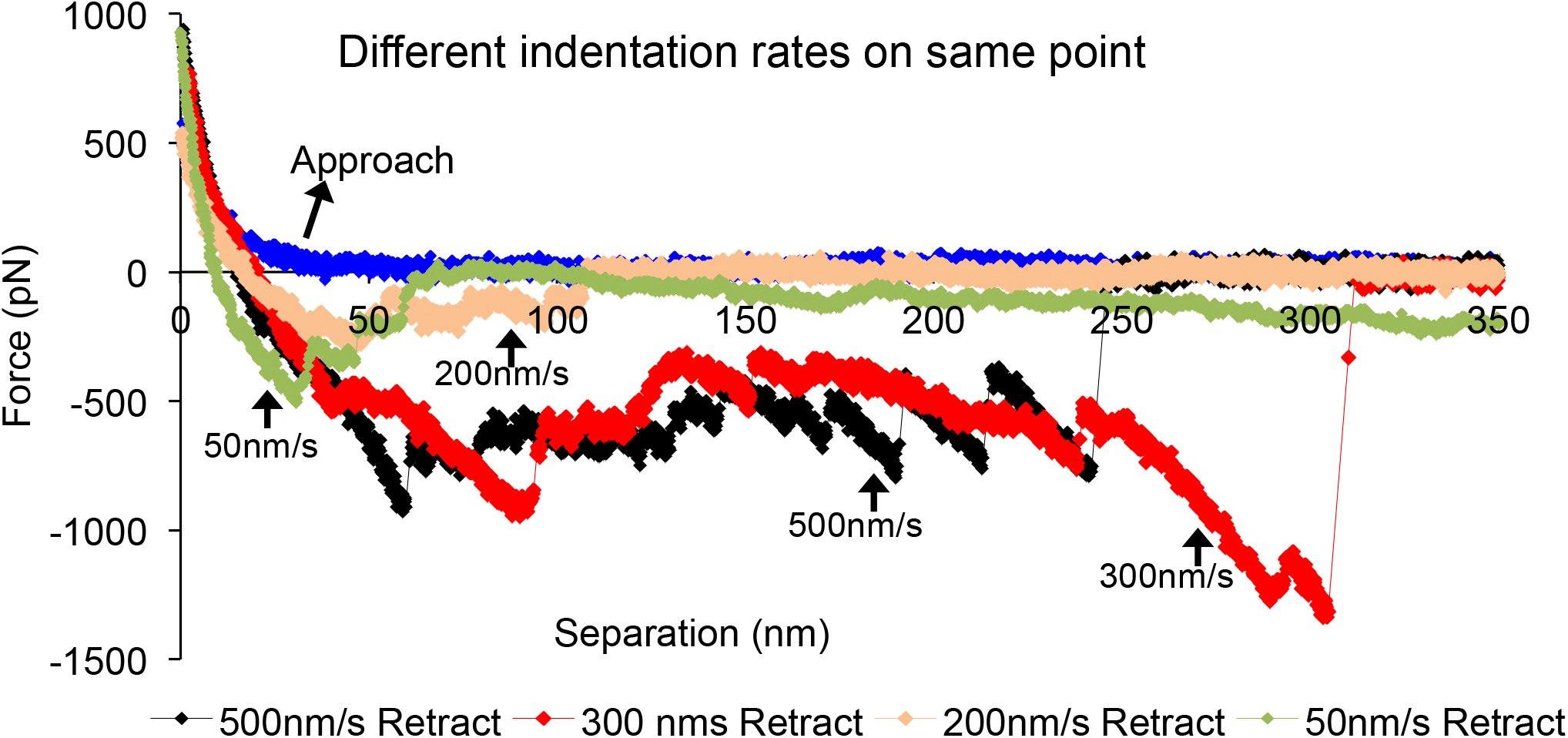
Retraction force curves observed at different indentation rates on the same location between the two virus glycan shields, indicating a rate-dependent change in adhesion patterns from man-man velcro-like at low rates to SA-SA slime-like at high rates.

Since there was no indication of mannose residues being exposed on the virus, the rate-dependent shift from SA- to mannose-like adhesions must involve rate-dependent access to the deeper mannose core or rate-dependent penetrability of the glycan shield. To test this premise, virus adsorbed on APTES surfaces was indented at different ramp rates with a bare AFM probe up to a maximum force of 1.2nN (Fig. 6A). At the higher ramp rates (towards 500 nm/sec), the resistance to indentation increased steadily in approach and decreased steadily with retraction (Fig 6B,C). Though there was hysteresis (i.e., the approach and retraction curves do not overlap due to permanent deformation occurring in the indented virus), an indentation force response that is proportional to the indentation strain indicates that the virus material is being compressed as a whole at these rates, without significant penetration of the material. However at lower indentation rates (towards ramp rates of 50 nm/sec), the force rise in approach was interspersed by plateaus and negative slopes (black arrows in Fig 6D,E). Sharp negative slopes typically indicate forced penetration of the material layers of the virus layers. The first penetration slope occurring at ~400 pN is likely to be in the soft glycan shield layer of the virus. Correspondingly, the retraction curves showed sharp negative forces (red arrow in Fig. 6E), which occurs if the bare tip is stuck at a penetrated layer and a build up in retraction force is required to release it. Figure 6F is a frequency plot of the incidents of negative slopes in virus approach force curves taken at different ramp rates. The higher rate approach curves were more likely to have 0 – 1 incidence of negative slopes (blue arrow), whereas the lower rates were more likely to 3 – 4 incidents of negative slope (orange arrow). Therefore a possible explanation for the rate dependent shift in glycan-glycan adhesion patterns is that the deeper mannose residues are being sampled at the lower rates where the glycan shield can be penetrated, whereas the surface SA residues are being sampled at the higher ramp rates where the glycan shield deforms as a whole. However, this explanation supposes that the surface SA residues are responsible for the slime-like adhesion and deeper mannose residues are responsible for the Velcro-like adhesion. We verify this supposition in the following experiment.

**Figure 6:**
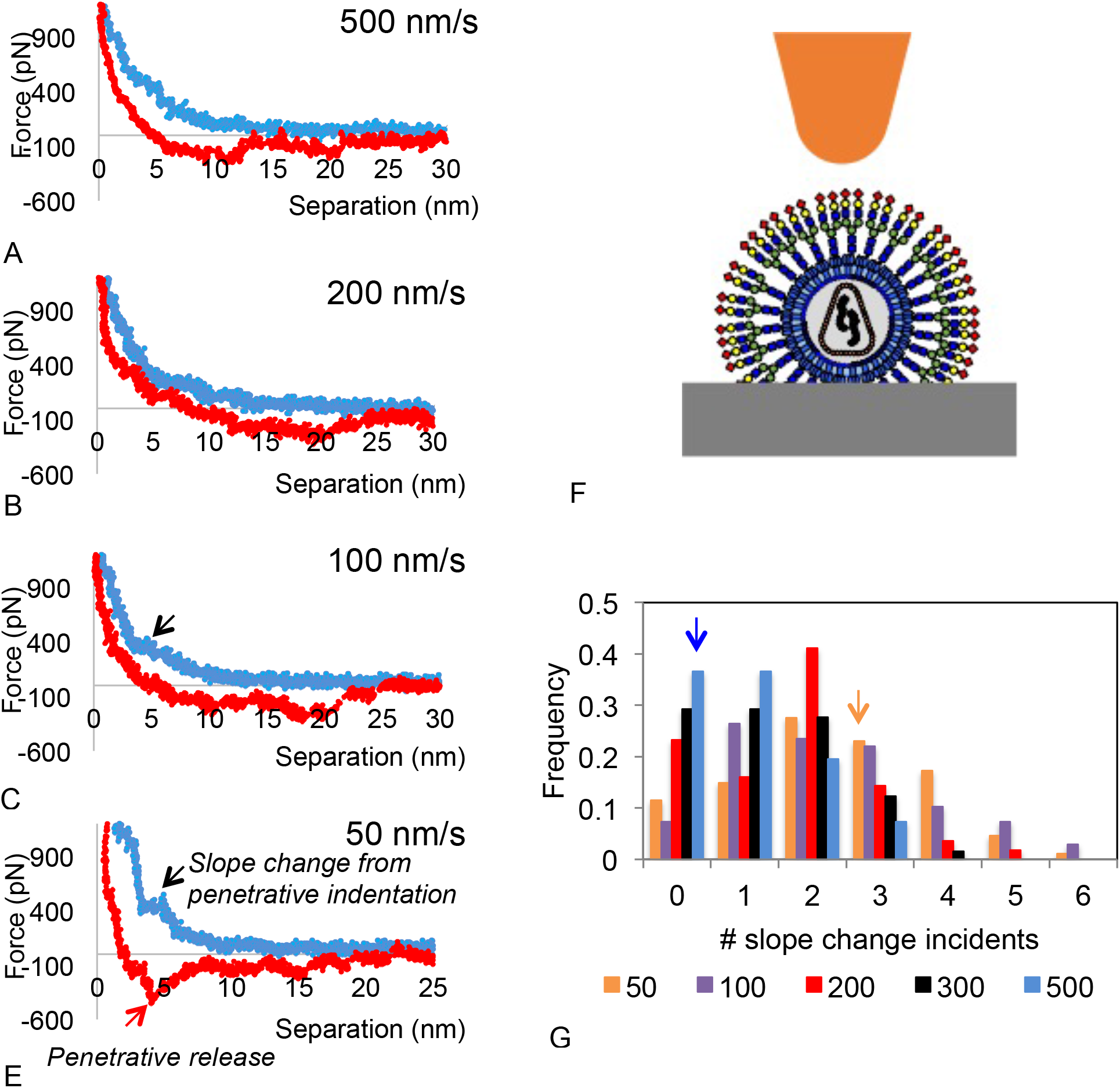
Rate dependence of the bare-probe indentation profile of a surface adsorbed virus: [A – E] Indentation force curve at different rates showing that as the rate decreases regions with negative slope appear during approach (black arrow in blue curve) and negative adhesion occurs during the retraction (black arrow in red curve). [F] Schematic of the indentation experiment. [G] Frequency plot of incidence of negative slopes in the approach curves, where 0 incidence is more common at higher rates (blue arrow) and 3 – 4 incidence is more frequent at the lower rates (orange arrow).

### 3.4 Slime- and Velcro-like self-adhesions localized at SA and mannose sugars, respectively

β-N-Acetylglucosaminidase (NAG) exposes the mannose core by cleaving the ß (1–2) glycosidic bonds between mannose and the GlcNAc residues above it (Fig. 7A). When two viruses with exposed mannose core were brought in contact and retracted, only Velcro-like adhesions were seen at low and high rates (Fig. 7B-D). The disappearance of the sawtooth negative forces confirms that the cleaved SA residues are the source of the slime-like adhesion. β-mannosidase enzyme was used to remove the mannose core by cleaving the β (1–4) glycosidic bonds between the mannose sugars and the GlcNAc residues below the core (Fig. 7E). When two viruses lacking mannose core but having exposed GlcNac residues were brought into contact and the retracted, there was no self-adhesion at all rates (Fig. 7F-H). The complete disappearance of velcro-like adhesions when the mannose core is removed validates the latter to be the source of these short-range adhesions between N-glycan shields (Fig. 6D).

**Figure 7.**
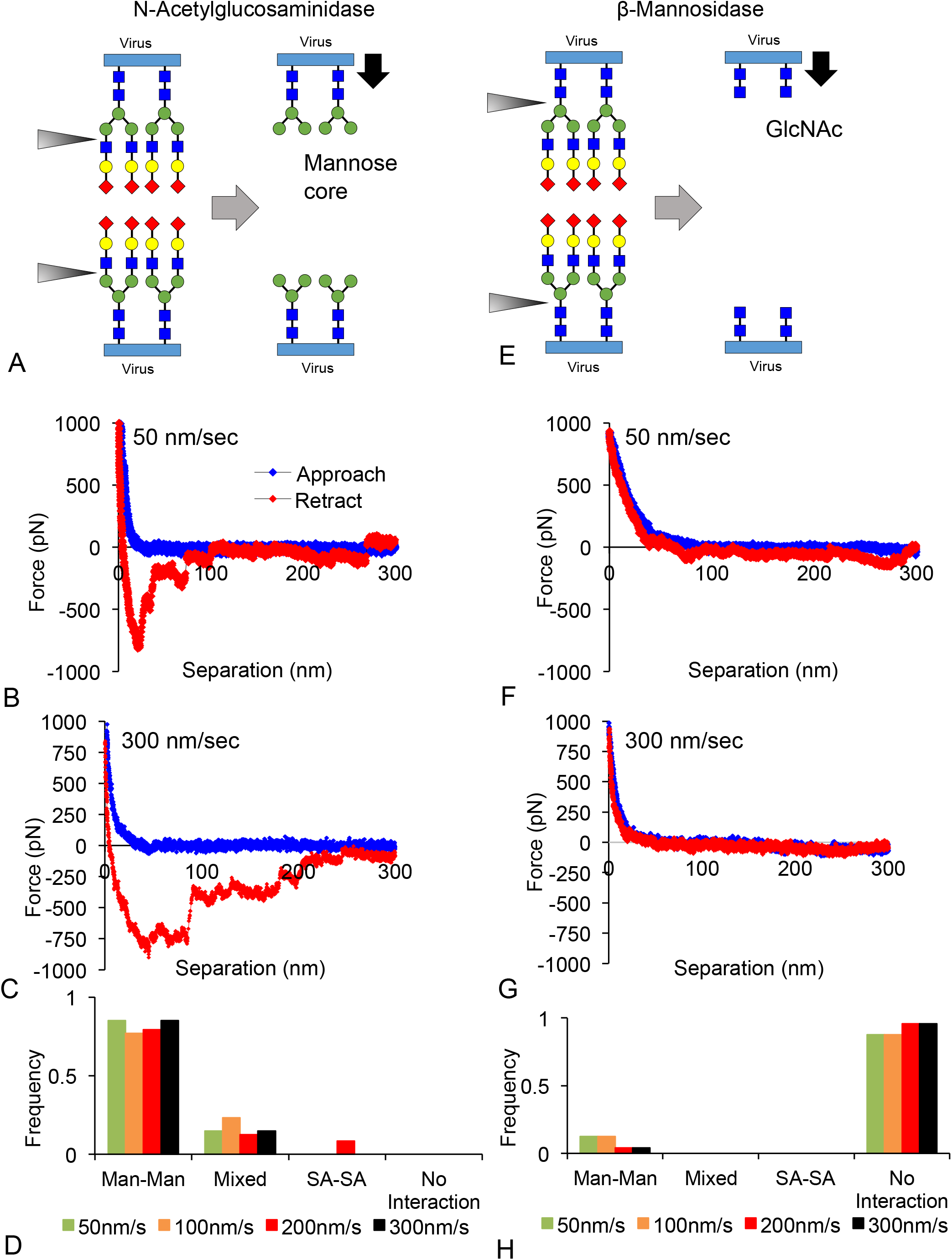
Nature of adhesive pull outs between two viruses when their mannose cores are exposed (right) and removed (left). [A] Complex N-glycan shield is cleaved with enzyme β-N-acetylglucosaminidase to expose the mannose core. [B] Representative indentation and retraction force curves between two viruses with exposed mannose core show velcro-like adhesions at lower ramp rates (~50nm/sec). [C] Representative FS force curves between two viruses with exposed mannose core exhibit velcro-like adhesions also at high ramp rates (~300nm/sec). [D] Distribution of adhesion types between two mannose cores as a function of ramp rate showing man-man velcro-like interactions occuring at all rates. [E] Complex N-glycan shield is cleaved with glycosidase β-mannosidase to remove mannose core and expose underlying GlcNAc residues. [F] Representative indentation and retraction force curves between two viruses with no mannose core and with exposed GlcNAc residues and showing no adhesion at low ramp rates (~50nm/sec). [F] Representative force curve between two GlcNAc exposed viruses showing no adhesion also at higher ramp rates (~300nm/sec) rates. [H] No self-adhesions were evident at all ramp rates between viruses with no mannose core and exposed GlcNAc residues.

The rate dependence of quantifiable features of the force response in both cases is shown in Supplemental Info. 2. The adhesion length and adhesion energy hovered around smaller values for all rates when the mannose core is exposed (which is consistent with man-man forces being short range), and became negligible when the mannose core is removed (consistent with mannose core being the source of the Velcro-like interactions). The indentation thickness was distributed similarly at all rates, confirming that comparable virus populations were being sampled. The indentation thicknesses for mannosidase-treated viruses (i.e., no mannose core) were however larger than the uncleaved (buried mannose core) and NAG-treated (exposed mannose core) virus, indicating a possible decrease in the structural stiffness of the virus following removal of the mannose core.

### 3.5 Velcro-like adhesion between mannose- and SA-terminated complex glycan shields

In earlier studies, no adhesions were observed between SA and mannose monolayers.(25) To understand how the monolayer findings with single sugars translated to multi-sugar glycan shields with terminal SA and mannose residues, we performed the following study. The virus on the AFM probe was cleaved with NAG to expose the mannose core (i.e., terminal mannose) and brought in contact with uncleaved virus (terminal SA) (Fig. 8A). Velcro-like adhesions were observed at all rates between the SA- and man-terminated complex glycan shield (Fig. 8B). A sample force curve is shown in Fig. 8C. The adhesion energy, for instance, distributed toward Velcro-like adhesion values at all rates (Fig. 8D). Velcro-like adhesions were also predominantly observed when the same position on terminal mannose- and SA-glycan shields was contacted at different rates (Fig. 8E). Even though the two glycan shields have terminal mannose and SA residues which are non-adhering in monolayers, penetrability and access to the mannose core appear to determine their net adhesion in FS. Frequency plots of the adhesion force and indentation thickness quantified at multiple rates are provided in Suppl. Info. 2.

**Figure 8:**
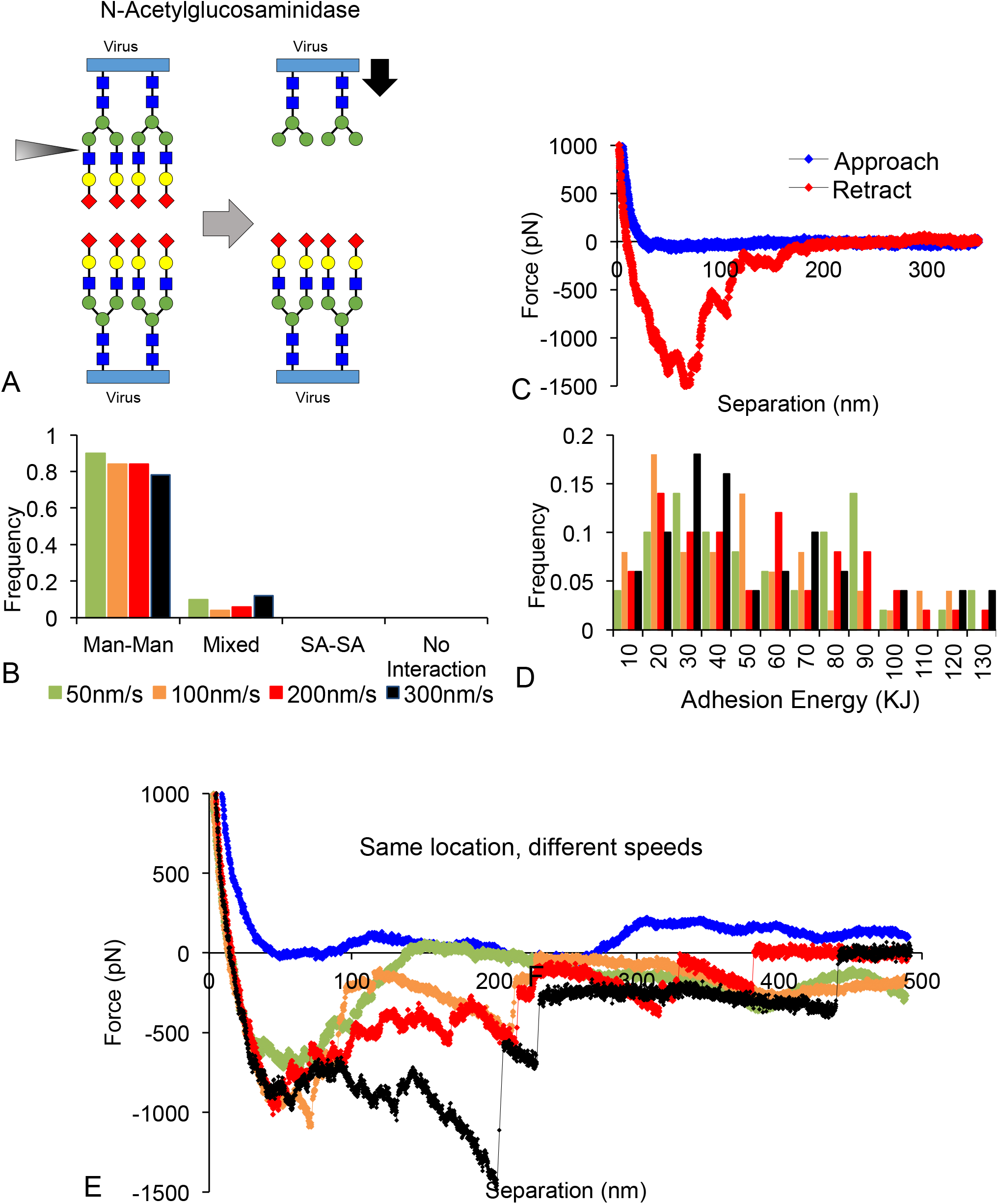
Cross adhesion between complex glycan shields that are SA- and man- terminated. [A] Virus attached to probe was cleaved by NAG to expose mannose core. [B] Representative interaction curve showing the man-man velcro-like interaction that occurs at all rates. [C] Frequency plot showing adhesions classified as man-man dominate at all rates. [D] Distribution of adhesion energy quantified from force spectroscopy curves showing similarly distribution at all rates (n > 20 for each ramp rates, 3 samples and 2 runs). [E] Change in adhesion profile at same point when indentation rates are changed; man-man adhesion releases were dominant at low rates whereas few mixed release occurred at higher rates in this case.

### 3.6 Virus self-aggregation in solution is determined by surface residues

We investigated how the self-adhesion patterns in force spectroscopy translated to self-aggregation in solution of viruses with different sugars of the N-glycan shield terminally exposed. Glycosidases were added to virus solution and virus aggregation was tracked after 1 hr incubation by changes in the DLS correlation curves. Lectins were added to verify the identity of the terminally exposed sugars. Virus aggregation and binding with lectins would increase the size of the diffusing species in solution (i.e., the hydrodynamic diameter D_H_) which is detected in DLS by a rightward shift of the virus correlation curve.

#### Sialic acid exposure (Fig. 9)

**Fig.9.**
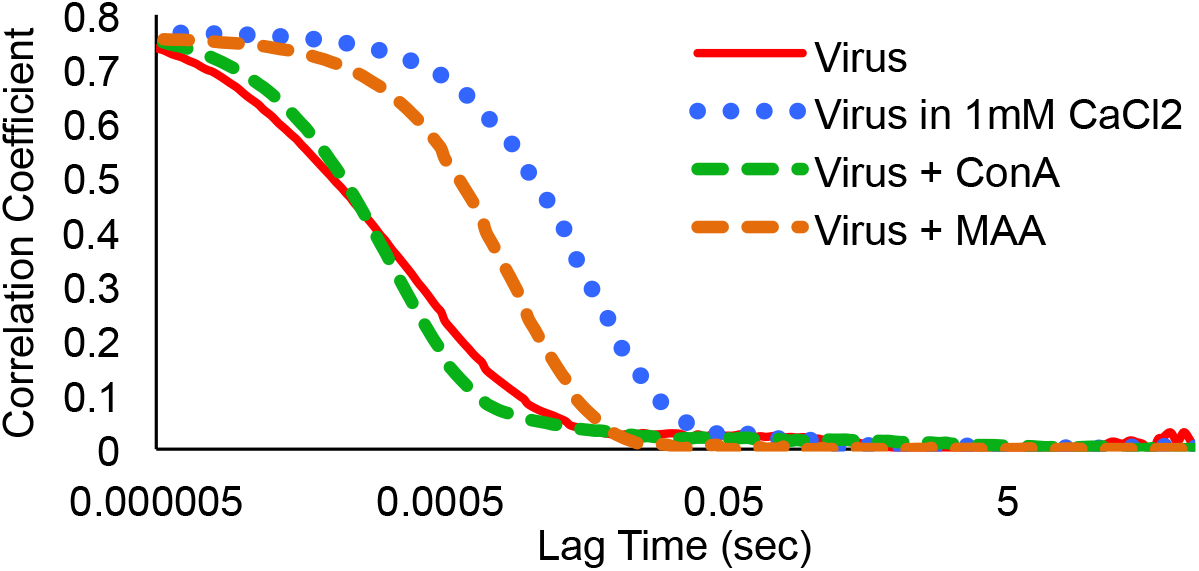
Verifying SA exposure on virus glycan shield by tracking shifts in the virus DLS correlation curve. The virus solution aggregated with SA lectin MAA (new exponential fall appears to the right in the virus correlation curve) and in the presence of Ca ion (virus correlation curve shifts to right), but not with mannose-lectin ConA (no rightward shift of correlation curve).

SA exposure on the virus was tested by the interaction with SA lectin MAA, mannose lectin ConA, and by aggregation with Ca^++^ ions. The hydrodynamic diameter of the virus is about ~200 nm in PBS. When MAA is added to the virus solution, the DLS correlation curve shifts to the right with the appearance of diffusing species in the range of 3μm, and indicating aggregation of the virus population. When Con A was added to virus solution, no rightward shift of the DLS curve indicating aggregation was observed. The lectin binding results confirm that the HIV-1 R-E-/VSV-G virus has terminal SA residues and insignificant terminal mannose. While surface-bound viruses exhibited slime-like self-adhesion in force spectroscopy, there was no corresponding self-aggregation between freely diffusing virus in solution. It is likely that virus self-aggregation in solution is prevented by the charge repulsion between surface SA residues. Accordingly when Ca^++^ ions, which screen charge repulsion, were added, large-scale self-aggregation of the virus occurred. The DLS correlation curve shifted rightward as aggregates of 2**μ**m D_H_ appeared in solution.

#### Galactose exposure (Fig. 10A)

**Fig. 10:**
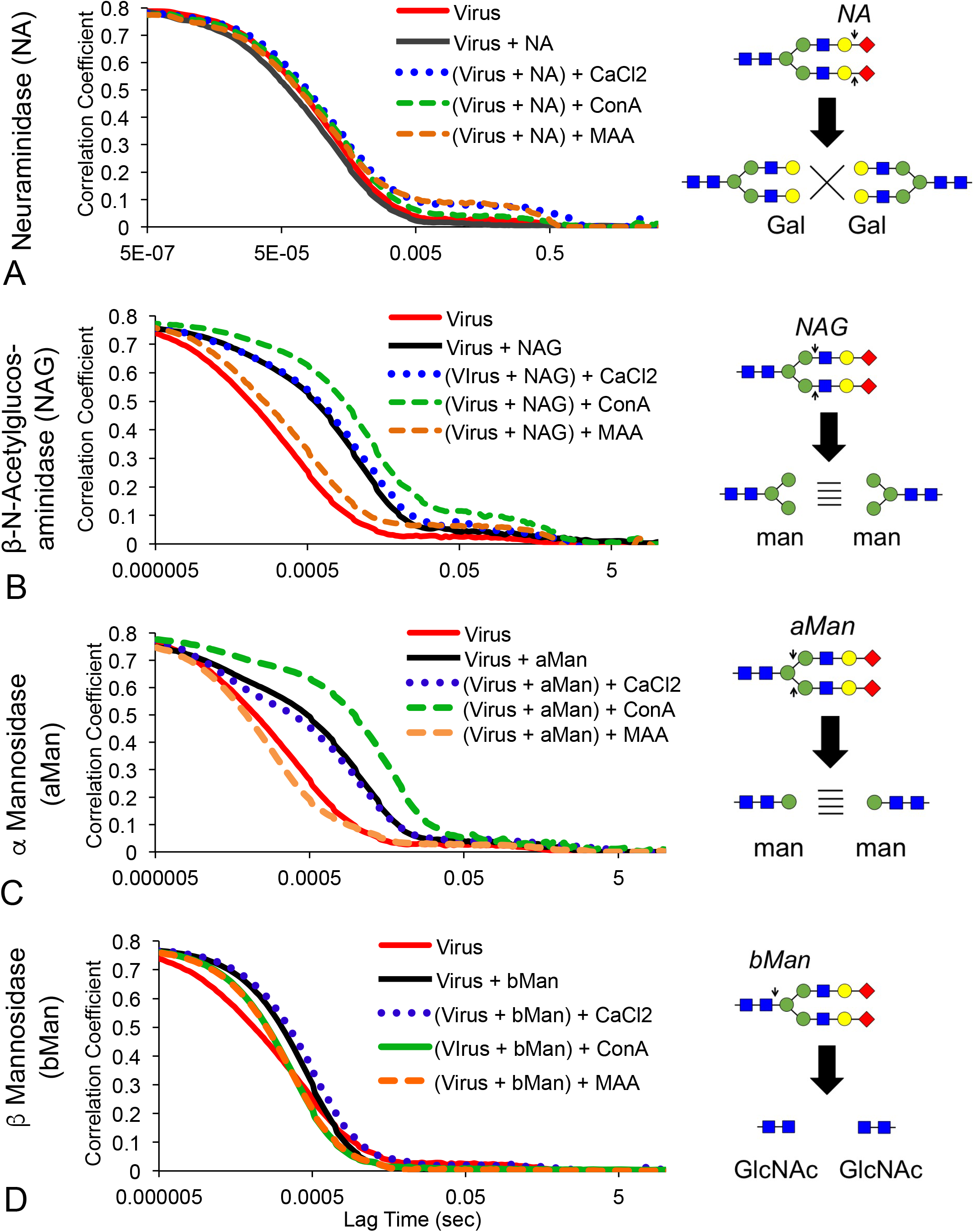
Self-aggregation propensity of each tier of complex N-glycan sugars in solution at 25C. As each tier of N-glycan sugar is exposed by glycosidases (shown on left), the DLS curve moves to the right if self aggregation of viruses with the exposed sugar tiers occurs and increases the size of the diffusing particles. Aggregation by SA lectin MAA and Ca^2+^ were also indicated by rightward shifts of the virus DLS curve and it indicates presence of terminal SA residues, whereas rightward shifts produced by ConA addition indicate presence of terminal mannose residues.

When Neuraminidase was added to virus solution to cleave SA and expose galactose residues, there was no rightward shift in the DLS curves even after 1 hour incubation, indicating absence of gal-gal self-aggregation. The complete removal of SA residues was confirmed by the absence of aggregation with SA-lectin MAA and Ca^++^ ions (no rightward shift). There was no aggregation with mannose-lectin Con A either. The absence of aggregation between Gal-terminated viruses is consistent with the force spectroscopy observation of no adhesion between two galactose monolayers.

#### Mannose exposure (Fig. 10B,C)

When the virus was cleaved with NAG to expose the mannose core, the hydrodynamic size increased from ~200 nm to ~500 nm (Fig. 10B). Removal of sugars is expected to decrease virus size, but the observed increase suggests that the viruses with exposed mannose core aggregates. Neither the SA-lectin MAA nor Ca ions aggregated NAG-cleaved virus, verifying there was no uncleaved terminal SA. Addition of ConA produced large aggregation, verifying that mannose residues are exposed. These results confirm that the Velcro-like self-adhesion observed between mannose cores in FS also manifest in solution. To check if the branched presentation of mannose in the N-glycan core is critical for Velcro-like self-adhesion, we removed the higher mannose residues with α-mannosidase (Fig. 10C). Again there was progressive aggregation, indicating that it is mannose residues that foster the self-aggregation and not necessarily the branched core architecture. Again, SA-lectin and Ca ions did not aggregate the α-mannosidase cleaved virus verifying complete removal of SA sugars, and mannose-lectin Con A aggregated the virus consistent with mannose exposure.

#### GlcNac exposure (Fig. 10D)

When β-mannosidase was added to the virus solution to remove the mannose core, no size increase corresponding to aggregation of viruses was observed. Cleavage of the distal residues was confirmed by the absence of aggregation with MAA, ConA, and Ca^2++^ ions. The lack of aggregation is consistent with force-spectroscopy findings in Fig. 7E-H where there were no self-adhesions between GlcNAc residues on surface bound virus.

The above results suggest that, unlike in force spectroscopy where penetrability of the glycan shield determined self-adhesion, only the terminal sugar determines self-aggregation of virus in solution. To further check this observation, we compared the self-aggregation of viruses with terminal GlcNAc residues that were either above the mannose core (produced by galactosidase cleavage) or below the mannose core (produced by β-mannosidase cleavage) (Fig. 11 A). Should glycan shield penetrability determine self-aggregation in solution, then the former with a mannose core beneath would exhibit different aggregation patterns than the latter which does not have a mannose core. The absence of self-aggregation in both cases, however, confirms that in solution the non-adhesive character of the terminal GlcNAc residues supersedes any penetrated adhesion from the mannose core. Figure 11B is a summary of the changes in D_H_ of the virus solutions as each tier of N-glycan sugars are exposed. Self-aggregation occurs (i.e., large jump in D_H_) only when mannose residues are exposed or when SA residues are in the presence of Ca^++^ ions.

**Fig. 11:**
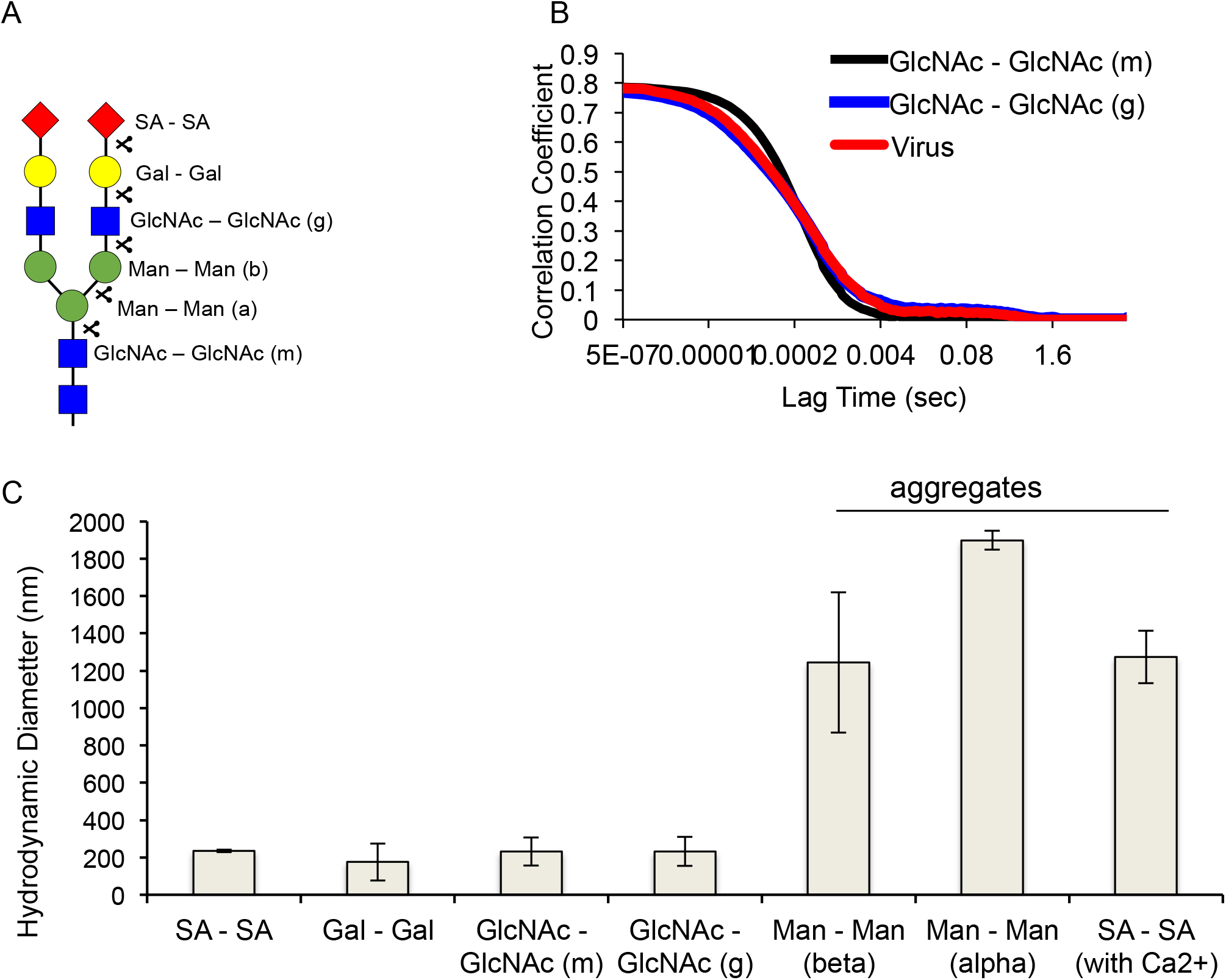
The self-aggregation pattern of N-glycan sugars in solution depends on the terminal sugar and not on penetrability of the glycan shield. [A] Summary of the different cleavages and therefore terminal sugars exposed in the solution. [B] DLS correlation curves comparing virus solutions with GlcNAc-terminal viruses having mannose core (g – generated by galactosidase cleavage) and GlcNAc-terminal viruses having no mannose core (m – generated by mannosidase cleavage). Both GlcNAc-terminal viruses do not show self-aggregation (i.e., no rightward shift of the DLS curves) indicating that penetrability and access to mannose core does not govern virus self-aggregation in solution. [C] Summary of the hydrodynamic diameters in solutions of viruses with different sugar terminals showing that only mannose and SA -terminated virus exhibit self-aggregation, but with the latter requiring Calcium ions.

## 4. Discussion

The goal of this study was to investigate how the placement of sugars in N-glycans affects the adhesion properties of biological surfaces. All N-glycans have a core of mannose residues, and complex type N-glycans has surface sialic acid residues. The complex-type N glycan shield of a HIV virus pseudotyped with VSV-G proteins was used as a biologically relevant platform for studying sugar interaction. Morphological and indentation characterizations of the HIV-1 R-E-/VSV-G pseudotyped virus had been detailed earlier.(25) In general the low indentation forces (< 2nN) observed in this study support the conclusions of the previous study that pseudotyping weakens the structural stiffness of the native HIV-1 virus. The complex N-glycan shields of viruses were bought into contact by force spectroscopy (FS) and by free diffusion in solution. Superficial sugars were cleaved using glycosidases to expose underlying sugars. Changes in FS self-adhesion and solution self-aggregation were tracked to understand the rules of sugar-sugar adhesion in glycan presentation.

### Slime-like and Velcro-like adhesions in N-glycan sugars

The interactions between glycan shields confirm previous observations with sugar monolayers that there are two types of self-adhesions among N-glycan sugars: (i) brittle or Velcro-like self-adhesions which release in single peaks close to the interface and originate from mannose residues; and (ii) tough or slime-like adhesions which release in multiple spurts over distances beyond the interface and originate from SA residues. (25) Removal of the respective sugars from the N-glycan shield led to loss of the adhesion types they foster.

### Glycan shield penetrability determines adhesion patterns in force spectroscopy

When both SA and mannose are presented in N-glycan architecture, the rate-dependent penetrability of the glycan shield appears to determine the type of self-adhesion manifested. At higher rates of indentation, the penetrability of the complex glycan shield decreases; a bare AFM tip compresses the glycan shield as a whole; and the slime-like self-adhesions from the surface SA residues dominate. At low rates, the penetrability of the glycan shield increases; a bare tip indents the shield with sharp rise and falls in force indicating penetration; and Velcro-like self-adhesions between buried mannose cores dominate.

It is also not clear what the source of the rate dependence in glycan penetration is. The rate dependence disappeared when SA and Gal residues were together removed from at least one of the N-glycan shields. When uncleaved (SA terminal) virus was brought in contact with NAG-cleaved (mannose core terminated) virus in FS, Velcro-like adhesions were observed all rates, indicating penetration of the uncleaved virus glycan shield and participation from the buried mannose at all sampled rates. The observation is consistent with previous reports where mannose monolayers on AFM probes penetrate complex N-glycan shields to elicit Velcro-like or brittle man-man adhesions.(25) There appears to be a general theme that mannose-terminal surfaces can penetrate SA-terminated glycan shields to latch on to the latter’s mannose core.

### Virus self-aggregation in solution is determined by the terminal sugar

SA- and mannose-terminal viruses self-aggregated in solution as was expected from their self-adhesion observed with FS. However there was a key difference in their aggregation patterns. Mannose-terminated glycans did not require Ca^++^ ions to self-aggregate whereas SA-terminated required Ca^++^ ions. So far we have observed mannose surfaces to self-adhere both in the presence and absence of ions,(24) in monolayer and glycan presentation.(25) However SA surfaces requires Na^+^ ions to adhere in FS, and Ca^++^ ions additionally to self-aggregate in solution.(25) The ion dependence supports previous conclusions (25) that the slime-like adhesions between SA residues are mediated by ions either to screen the charge repulsion or to bridge the interaction between the two negatively charged surfaces.

The other two N-glycan sugar tiers, galactose and GlcNAc, did not promote self-aggregation in solution. The solution behavior is in general agreement with FS observation on these single sugars. Monolayer studies in FS had reported that galactose sugars do not self-adhere. (24) N-glycan shields with only GlcNAc residues did not show self-adhere in FS (Fig. 7E-H). It appears that the terminal sugar alone determines the self-aggregation pattern of viruses in solution. GlcNAc terminated viruses did not self-aggregate in solution irrespective of whether a mannose core was present beneath the GlcNAc layer or not. In other words, penetrable access to the mannose core did not determine self-aggregation patterns in freely diffusing virus in solution. We do note that the study did not explore how ramp-rate dependence of penetration observed in FS translates to shear-rate dependence of aggregation in solution.

### Rules of interaction applicable beyond N-glycan presentation of sugars

Virus with exposed mannose core self-aggregated even when the first tier of the mannose core was cleaved (Fig. 10C). The shortrange self-adhesion between mannose residues had been observed when the sugar presentation is disordered in monolayers. In other words, mannose presentation in N-glycan architecture appears not critical for Velcro-like self-adhesion to occur, and the findings could be translatable to other systems where mannose is abundance. The endosperm of seeds (dates, coffee beans, ivory nuts, etc.) is rich in polymers of mannose called mannans.(29) These polymers are tightly packing since they serve as a dense store of energy for the seeds. X-Ray crystallography reveals organized hydrogen-bonding across stacked mannose faces responsible for the tightly packed in mannose polymers. (30–32) Similar shortrange hydrogen bonding interactions could be responsible for the Velcro-like adhesions between mannose residues in glycans. Interestingly the tight packing among mannose polymers is loosened in mucilaginous materials (guar gum, locust bean gum, cassia gum) by interspersing the polymer with galactose. (29,33) It is known that greater the galactose:mannose ratio, higher the swelling or loose-packing of the galactomannan polymers.(34) This observation in galactomannan polymers is in line with our findings that galactose sugars do not adhere to other galactose residues or mannose residues, and therefore would interfere with the tight-packing of mannose polymers.

### Biomedical and Bioengineering implications

The consistency of the sugar self-adhesion findings between monolayer-monolayer, monolayer-glycan shield, and glycan shield-glycan shield studies suggests that the precise presentation of the sugars is not as critical a factor for sugar-sugar adhesions as it is for protein-sugar binding. In the case of the latter, the presentation of the sugar determines how it fits within the binding sites on lectins. This implies that the rules of sugar interactions determined with N-glycan shields could be extended to engineering sugars systems and monolayer coatings where the presentation of sugars is not as defined. For instance, these rules would help understand how mannose-coated nanoparticles, for instance, distribute among the glycosylated surfaces in the host.(35,36) Moreover, the composition of glycan changes with age, disease, stress, and environment factors.(37) SA and mannose sugars, which influence how neurons interconnect, are diminished in neural disorders such as multiple sclerosis (38–40); distal SA decreases during aging but distal galactose increases; mannose expression increases in breast cancer (41) and SA expression is altered in cancer. (42) Environmental stresses also change the N-glycan composition.(43–45) Establishing guiding rules for how changes in sugar composition alter the interfacial environment will provide a new perspective for interpreting the biological changes occurring in these conditions. Our findings also suggest that the interfacial properties of biological systems will change drastically when the composition of surface sugars change.

---

**Supporting Information 1:**
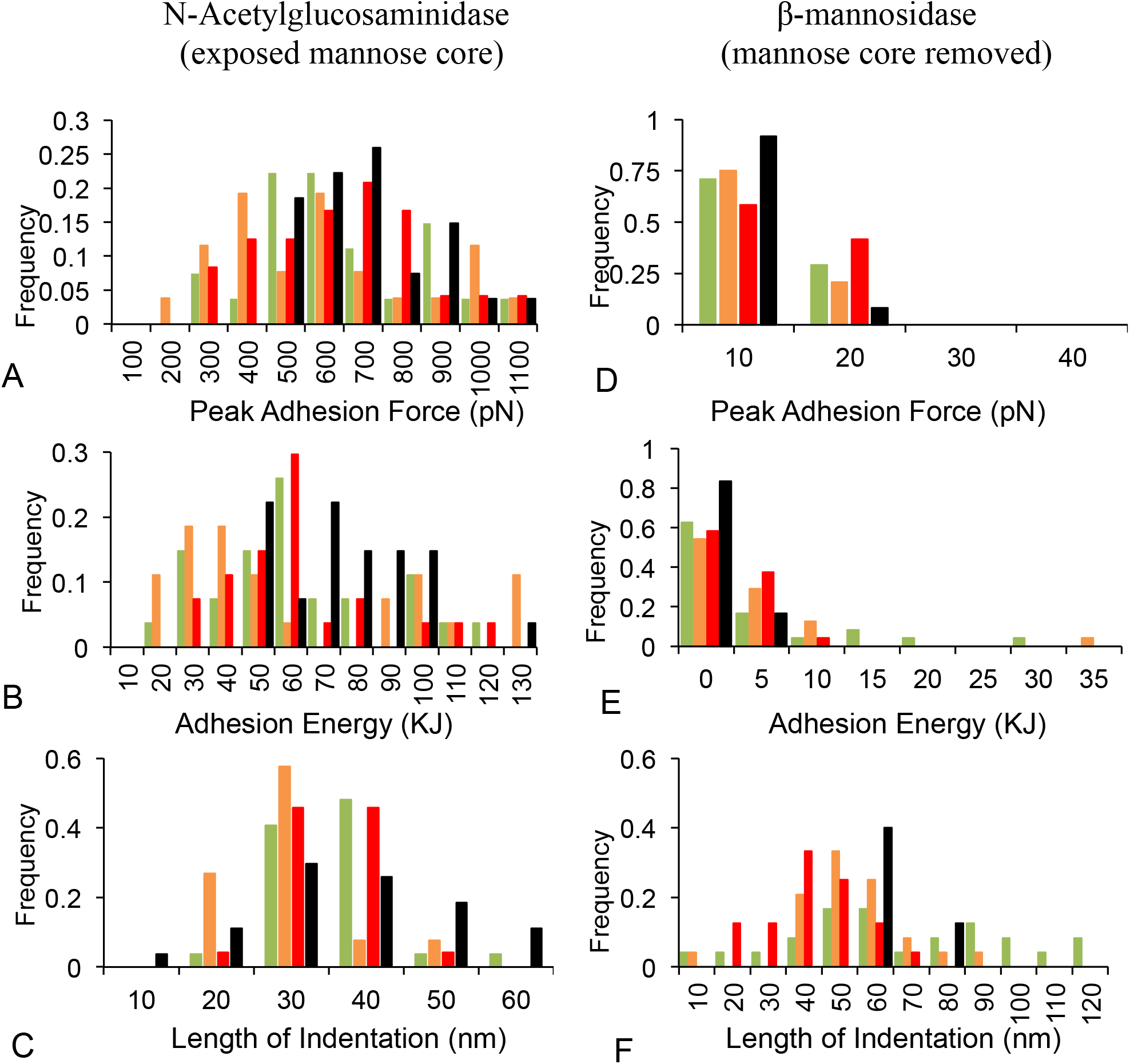

Distribution of adhesion parameters when two complex glycan shields cleaved to expose mannose cores are brought in contact (A,B,C) and when the mannose core is removed from both glycan shields (D,E,F)

**Supporting Information 2:**
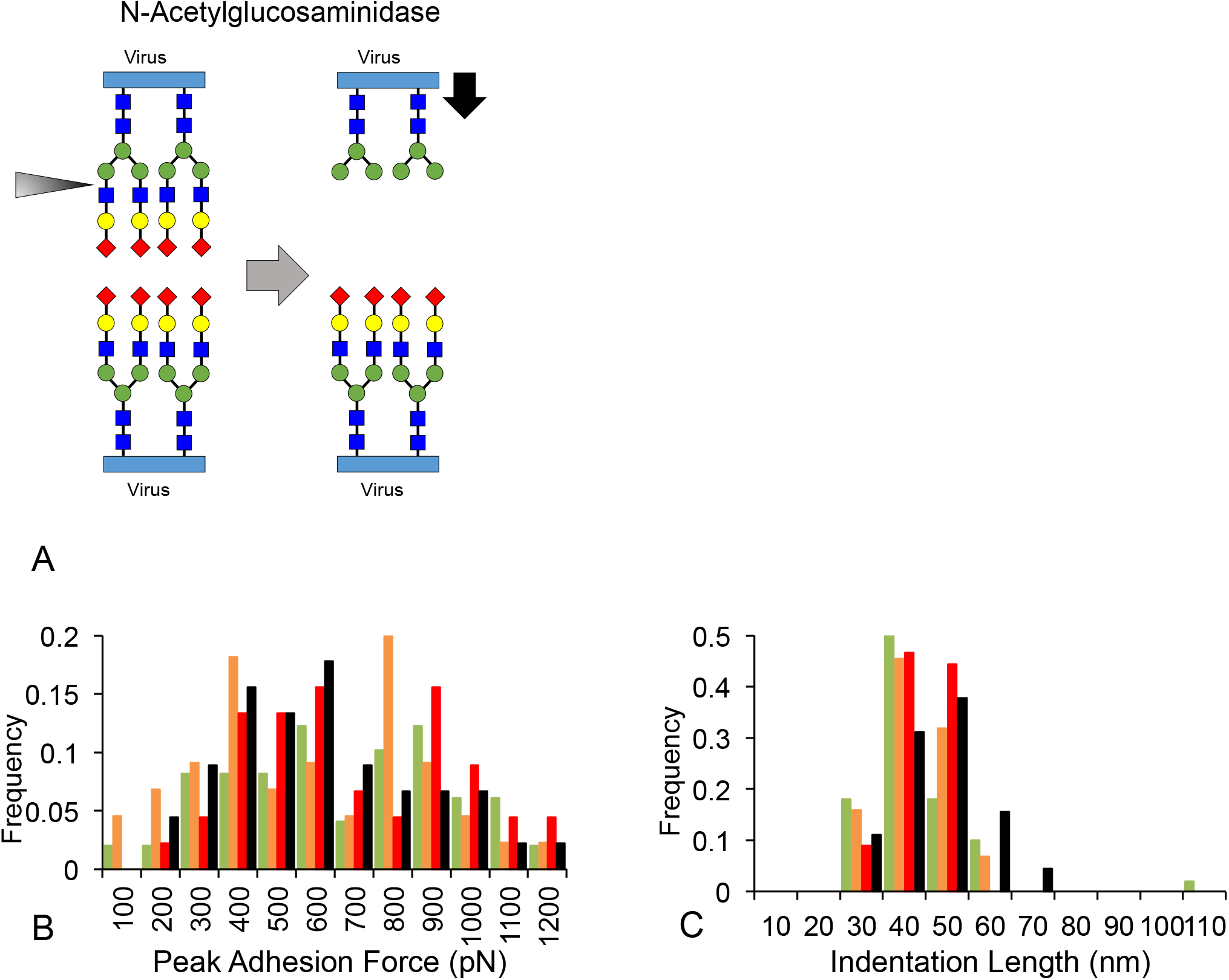

Distribution of peak adhesion forces (B) and indentation lengths (C) when an intact glycan shield is brought in contact with the mannose core exposed on another glycan shield (A).

## ACKNOWLEDGEMENTS

We are also thankful to Dr. J.W. Mitchell for access to the Nanomaterials lab at Howard University, and to Mr. James Griffin. We thank Drs. Anna Allen and Dr. Saswati Basu for reviewing the manuscript. EO was supported by the Just Julian Fellowship from Howard University. This project was funded in part with federal funds (UL1TR000101 previously UL1RR031975) from the National Center for Advancing Translational Sciences (NCATS), National Institutes of Health, through the Clinical and Translational Science Awards Program (CTSA), a trademark of DHHS, part of the Roadmap Initiative, “Re-Engineering the Clinical Research Enterprise.”

